# A ventral striatal learning signal reflecting individual differences in the success of fear extinction

**DOI:** 10.1101/2025.08.26.672387

**Authors:** Elena Andres, Hu Chuan-Peng, Anna M. V. Gerlicher, Oliver Tüscher, Raffael Kalisch

## Abstract

Individual differences in fear extinction learning are centrally involved in anxiety vulnerability. We here investigate individual extinction differences using a model-free, data-driven approach, by applying Latent Class Growth Modeling (LCGM) to four in-house data sets from altogether N=234 healthy male participants. This revealed two distinct trajectory classes: fast extinguishers and slow extinguishers. This pattern was replicated in two independent public data sets (total N=275, female and male). In a subset of the in-house samples with functional magnetic resonance imaging (fMRI) data (n=122 males), we investigated the neural correlates of class membership, focusing on the ventral striatum (VS), a key area previously implicated in encoding extinction prediction errors (EPE). We found that fast extinguishers exhibited VS activity at the time of unconditioned stimulus omission early in extinction, consistent with an EPE signal, whereas this signal only appeared late in extinction in slow extinguishers. These findings suggest that extinction success is shaped by how the VS learns safety.

## Introduction

Successful extinction learning is a key protective factor against the development of anxiety and stress-related disorders following trauma exposure (1). Conversely, impaired extinction has been consistently observed in patients with anxiety disorders when compared to healthy controls (2–4). These findings highlight the importance of extinction mechanisms in the etiology of anxiety disorders and motivate a focus on understanding individual differences in extinction learning even within healthy populations (5).

When a bad outcome that we expect does not materialize, we experience surprise and relief. Fear extinction learning is thought to be driven by this unexpected omission of the feared outcome. In extinction, a stimulus (conditioned stimulus, CS) paired during fear conditioning with an aversive stimulus (unconditioned stimulus, US), such as pain, is repeatedly presented without pairing with the US, and conditioned fear responses (CRs) towards the CS eventually decline (6). Classical associative theories like the Rescorla-Wagner model (7) postulate that the magnitude of the CR over the course of conditioning and extinction reflects the strength of the CS-US association, understood as prediction of the occurrence of the US conferred by the CS. For each new CS trial, the current US prediction (determining the aversive value of the CS and, hence, CR magnitude) is updated commensurate with any expectancy violation, or prediction error (PE), that may have occurred at the last trial. If the occurrence of the US is unexpected, such as in an initial conditioning trial, the discrepancy between US occurrence (1) and US prediction (0) leads to a corresponding increase of US prediction and CR in the next trial. If, in extinction, the US does unexpectedly not occur, this inverse PE reduces US prediction and CR in a subsequent trial. In both experimental phases, the unsigned amplitude of the PE itself decreases to the extent that US occurrence (conditioning) or non-occurrence (extinction) have become expected.

PE-based extinction learning shows strong analogy to appetitive learning (8), as unexpected US omission like unexpected reward are better-than-expected outcomes and are both experienced as pleasant. It has therefore been proposed that extinction learning is mediated by the same mesolimbic dopamine (DA) system that mediates reward learning (9,10). Specifically, it has been hypothesized that the extinction prediction error (EPE) should not be encoded neurally like an inverse aversive PE signal but like a dopaminergic reward prediction error (RPE) signal. In agreement with this proposal, animal studies have observed phasic activity of ventral tegmental area (VTA) DA neurons during extinction, specifically at the time of US omission, that conforms to theoretical expectations: high in early extinction trials and decreasing over the course of extinction, in correlation with the decrease of the CR; further, the signal could be shown to be necessary for extinction (11–14). Also, in agreement with an identity of EPE and RPE, the nucleus accumbens has been identified as the likely projection area of these neurons (12,14). In humans, computational functional magnetic resonance imaging (fMRI) studies have demonstrated a ventral striatal signal at US omission following a trial-by-trial time course predicted by the Rescorla-Wagner model (10,15). The appetitive theory of extinction (10) assumes that activation of the appetitive system during fear extinction may lead to CR reduction via antagonistic inhibition of the aversive system, as postulated in opponent-systems theory (16–18).

Applying associative models to learning data involves estimating the trial-wise PE signal from the trial-wise CR time course, such that – in the case of extinction learning – the observed decline of the CR is reflected in an initial, positively signed EPE that subsequently also declines in amplitude (10,15). For the purpose of time course fitting, the Rescorla-Wagner model as well as its later variants (19) all employ constants (learning rate α) that are multiplied with the PE term (occurrence minus prediction) to express to what extent the PE signal is translated into an update of the US prediction and, hence, can explain CR reduction. Higher learning rates imply quicker updating and faster CR reduction (faster extinction). Learning rates can be estimated individually. This allows for quantifying individual differences in the speed, or success, of extinction. For instance, Raczka et al. (10) have found evidence that a variant of the DA transporter (DAT) gene is associated with relatively higher learning rates (that is, faster extinction).

Because the learning rate and the PE signal are multiplicatively combined in a common updating term, the associative modeling approach does not permit to tell the possibility that faster extinguishers produce (initially) higher EPEs than slow extinguishers from the possibility that they generate similar EPE signals as slow extinguishers but translate these signals more efficiently into a reduction of the CR. The latter may be because they more easily update their CS-based US predictions following an EPE and/or because their updated predictions more quickly lead to CR inhibition. (Considering the likely appetitive nature of extinction learning, this latter possibility would imply a faster build-up of reward predictions and/or a better antagonistic inhibition of the aversive system.). Another limitation of associative models is that they all assume a strictly monotonous (gradual) decline of CRs during extinction that is commensurate with the development of the EPE. As such, they find it hard to incorporate the possibility that learning may occur in a more abrupt or step-wise fashion. Step-wise learning may occur when PEs need to accumulate up to a certain threshold to entail updating (for instance because of very strong prior expectations about the CS-US association) or when updated associations need to pass a threshold to entail CR reduction. Such relationships are difficult to model unless one introduces strong assumptions. Step-wise learning may also be caused by sudden insights or shifts in contingency beliefs, better accommodated by explicit reasoning- or model-based learning models (e.g.(20)). For the study of individual differences, this additional limitation of associative modeling implies that the approach may not capture between-subject differences that lie in the shape, rather than the speed, of learning.

In the present study, we abandoned the associative modeling framework in favor of a purely data-driven approach towards identifying individual differences in extinction learning and its underlying neural substrates.

There has been substantial effort to characterize the heterogeneity of extinction learning trajectories in rodents (21–25) and humans (26–29). Two studies (22,27) used latent class growth modeling (LCGM) of trial-wise responses, a data-driven method that is specifically tailored for identifying subgroups with distinguishable temporal profiles. In humans, LCGM analyses of subjective ratings of fearfulness and US expectancy have revealed that slow extinction learning is associated with increased state and trait anxiety and higher likelihood of anxiety-related disorders (26,28). Furthermore, a large-scale study applied LCGM to trial-wise skin conductance responses (SCRs) and identified distinct learning subgroups highlighting heterogeneity in fear response trajectories (27). These studies suggest that LCGM of human CR time courses during extinction is suitable to detect meaningful individual differences in extinction learning.

We thus applied LCGM to trial-wise SCR data from four studies conducted in our lab in healthy normal participants in which differential fear conditioning and extinction were conducted following an identical protocol (on day 1 and day 2, respectively). To assess the generalizability of this approach, we additionally applied the same LCGM procedure to two publicly available open SCR datasets (30,31). Unlike in other work from our group (10,15), the four in-house experiments were not designed, and not optimized, for associative modeling.

Having identified subgroups of extinction responders in these data, we then asked if we could identify differences in extinction-related neural activity between these groups in the subset of three studies performed in the MRI scanner. We tested a single hypothesis, namely that differences in extinction learning would be related to differences in how the ventral striatum (VS) reacts to US omission at the offset of the CS that was previously paired during conditioning with the US (CS+). The CS+ offset is the time point at which an EPE is registered, and the VS is the area that would be expected to show fMRI signal changes conformal with EPE encoding if the appetitive, DA-based account of extinction learning were right (9). We thereby tried to gain evidence for an involvement of ventral striatal EPE signals in individual differences in extinction learning unconfounded by potential individual differences in US prediction updating or CR inhibition and not restricted to the monotony assumptions of formal associative learning models.

## Methods

### Datasets

Data used in this paper contains four datasets from four different experiments using the identical fear conditioning (day 1) and extinction (day 2) paradigm. Data has been published for dataset 1 (32), dataset 2 (33), and dataset 3 (34). Dataset 4 has not yet been published separately. The purpose of these experiments was to study effects of L-DOPA on either extinction learning (dataset 4, L-DOPA administration before day 2 paradigm) or extinction consolidation (datasets (1-3), L-DOPA administration after day 2 paradigm). As class analysis was performed on day 1 and day 2 data, we excluded data from participants in dataset 4 who received L-DOPA before the day 2 experiment to avoid any confounding effects on the classification. Furthermore, day 3 data was left out in the LCGM analysis. An overview of the paradigms is shown in Fig. 1.

**Figure 1.**
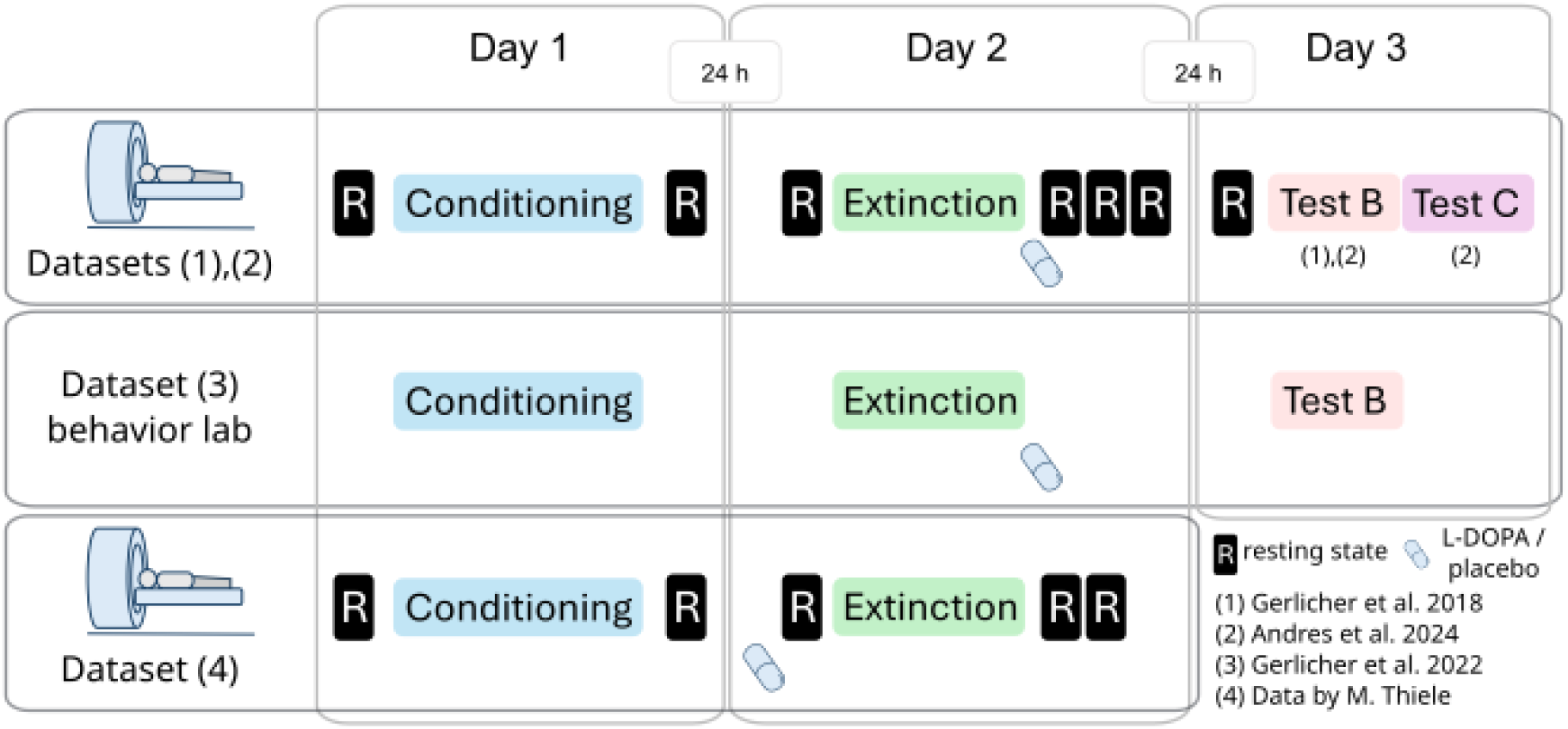
Overview of experimental design. Day 1: During fear conditioning on day 1, participants were presented with two geometrical figures serving as CSs in a context A. The CS+ was followed by painful stimulation (US) in 50% of the trials, the CS-never was reinforced and served as a control stimulus (CS-). Day 2: During extinction on day 2, participants were presented with both CSs again in another context B. However, the CS+ was not reinforced with a US anymore. Immediately before (dataset 4) or after extinction learning (datasets 1-3), participants were administered either a placebo or a L-DOPA pill. Day 3: At an extinction memory test on day 3 (datasets 1-3), participants were presented with both CSs in the absence of the US again in context B (the context in which the extinction learning took place). Dataset 2 further included a test in a new context C, which is not analyzed here. Datasets 1, 2, and 4 include fMRI data from all phases. The resting-state fMRI scans (R) in datasets 1, 2, and 4 are not relevant for the current analyses.

### Participants

Recruitment was restricted to male participants, since the estrous cycle interacts with extinction memory consolidation (35–39) and DA can have opposing effects on extinction depending on estrous cycle phase (40). A board-certified physician screened participants for contraindications of L-DOPA intake, current physiological, neurological or psychiatric disorders, excessive consumption of nicotine (>10 cigarettes/day), alcohol (>15 glasses of beer/wine per week) or cannabis (>1 joint/month), participation in other pharmacological studies and skin conductance non-responding (assessed by eSense Skin response, Mindfield® Biosystems Ltd., Berlin, Germany). Drug abuse was assessed via a urine test (M-10/3DT; Diagnostik Nord, Schwerin, Germany). The experiment was approved by the local ethics committee (Ethikkommission der Landesärztekammer, Rhineland-Palatinate, Germany) and was conducted in accordance with the Declaration of Helsinki.

### Experimental Design

All experiments consisted of the same paradigm on day 1 and day 2. Whereas experiments 1, 2, and 4 were implemented in the MRI scanner, experiment 3 was done in a behavioral lab. **Experiments 1-3**: For a detailed protocol see methods in (32–34). Briefly, a three-day fear conditioning and extinction paradigm was implemented. The experiment consisted of a fear conditioning phase on day 1 in context A, an extinction session on day 2 in context B and a test on extinction memory retrieval on day 3 in the original extinction context (B) and in a new context (C, only experiment 2, not considered in this work). **Experiment 4**: In this experiment, only the day 1 and day 2 paradigms were implemented. For experimental design see Fig. 1.

### Stimuli

Two black geometric symbols (a square and a rhombus) presented in the center of the screen served as CSs. The symbols were super-imposed on background pictures of one of three different grayscale photos (living room, kitchen, and sleeping room), which served as contexts A, B, and C. The assignment of symbols to the CS+ and CS- and background to the conditioning, extinction, or context C were randomized between participants and groups. In order to diminish the risk of low visual feature differences between the CS+ and the CS-, the contrast and luminance were adjusted between stimuli using SHINE toolbox (41). Stimuli were presented using Presentation Software (Presentation®, Neurobehavioral Systems, Inc., Berkeley, CA, USA). A painful electrical stimulation consisting of three square-wave pulses of 2 ms (50 ms inter-stimulus interval) was employed as US. Pain stimuli were generated by using a DS7A electrical stimulator (Digitimer, Weybridge) and delivered on the right ankle through a surface electrode with a platinum pin (Specialty Developments, Bexley, UK).

### Experimental Procedure

Day 1 - Fear learning. Upon arrival for testing, participants filled out the state anxiety questionnaire of the state-trait anxiety inventory (STAI-S (42). Participants were placed in the MRI scanner (experiments 1, 2, 4) or in the behavior lab (experiment 3) and SCR and pain stimulation electrodes were attached. Before the start of the experiment participants were familiarized with the experiment. Familiarization consisted of two CS presentations each in all possible contexts and practical training of fear and US-expectancy ratings. US-intensity was then calibrated to a level rated as “maximally painful, but still tolerable”. Participants were instructed that the experiment was distributed across two (experiment 4) or three days, that one symbol would never be followed by an electric shock and that their task was to find out what rule applied to the other symbol. The paradigm started and ended every day with US-expectancy ratings for each CS. Afterwards, a picture of the context appeared on the screen. The context picture remained on the screen continuously throughout the given experimental phase. CSs were presented for 4.5 seconds. In case of reinforced CS+ presentations, USs were delivered such as that they co-terminated with CS presentations. Inter-trial intervals (ITI) lasted 17, 18, or 19 seconds. Trial order in all experiments was randomized in such a way that not more than two trials of the same type (i.e., CS+, CS-) succeeded each other. During conditioning (Fig. 1) in context A on day 1 participants were presented with 10 CS+ and 10 CS-trials. Notably, 5 out of 10 CS+ presentations (i.e. 50%) were reinforced. Conditioning lasted for approximately eight minutes. An eight-minute resting-state fMRI scan was conducted before the familiarization and immediately after the experiment. Subsequently, electrodes were detached and participants filled out a list of questions designed to assess contingency knowledge.

Day 2 - Extinction learning and consolidation. Approximately 24 h (±2 h) after the start of the day 1 experiment, participants returned to the laboratory. After filling out the STAI-S, participants underwent the extinction training in the scanner (experiments 1, 2). After attaching the electrodes, participants were instructed that the experiment would continue, and that their individual US strength from day 1 was applied. During extinction, participants were presented 15 CS+ and CS-trials in context B, approximately lasting twelve minutes. There were no US presentations. Subsequently, participants were taken out of the scanner briefly to detach the electrodes and give participants either a placebo or an L-DOPA pill. After oral pill intake, participants stayed under observation for 90 minutes. Eight-minute resting-state scans were performed immediately before the experiment and at ∼5, 45, and 90 minutes after pill intake. Before leaving the laboratory, participants filled out the STAI-S and a questionnaire designed to assess possible side-effects of L-DOPA intake. Experiment 3 again was implemented in the behavior lab without any scans. L-DOPA or placebo was administered after extinction learning. In experiment 4, participants received the L-DOPA or placebo pill at arrival. Forty-five minutes later the same procedure including one resting-state before and two after extinction learning was implemented. Experiment 4 finished after day 2.

Day 3 - Test. Experiment 1-3: Approximately 24 h (±2 h) later, participants returned and filled out the side-effects questionnaire and the STAI-S. After electrode attachment, participants were instructed that their US strength from day 1 was applied and that the experiment would continue. Participants were then presented 10 CS+/10 CS-in context B first (approximately eight minutes) and subsequently the same number in context C (experiment 2, not considered in this work, see Fig. 1). An eight-minute resting-state scan was collected immediately before the start of the experiment on day 3 in experiments 1 and 2. Experiment 3 again was implemented in the behavior lab.

### Drug Treatment

Participants were randomly assigned to the L-DOPA or the placebo group, with the restriction that groups were matched on STAI-T (42) scores by a person not involved in the experiments. Participants were administered either 150/37,5 mg L-DOPA-benserazide (Levodopa-Benserazid-ratiopharm®, Germany; for dosage see (43,44)) or a visually identical capsule filled with mannitol and aerosol (i.e. placebo). Drugs were prepared and provided by the pharmacy of the University Medical Center Mainz and administered in a double-blind fashion.

### Skin Conductance Responses (SCR)

Conditioned fear responses were assessed by CS elicited SCRs. Electrodermal activity was recorded from the thenar and hypo-thenar of the non-dominant hand using self-adhesive Ag/AgACl electrodes prefilled with an isotonic electrolyte medium (EL-507, BIOPAC® Systems Inc., Goleta, California, USA) and the Biopac MP150 with EDA100C device. The raw signal was amplified, and low pass filtered with a cut-off frequency of 1 Hz. The onset of SCRs was visually scored offline in a time window from 900 to 4000 ms after CS onset. The amplitude of SCRs was then calculated by subtracting the onset SCL from the subsequent peak, using a custom-made analysis script. The SCRs with amplitudes smaller than 0.02 μs were scored as zero and remained in the analysis. Participants were excluded if they had zero valid SC responses on one of the three days. To normalize distributions, data was log-transformed (+1 and log) and range-corrected for each participant and experimental session (i.e. (SCR - SCRmin) / SCRmax (45)).

### US-expectancy ratings

Participants were asked to indicate the expectancy of receiving an electric pain stimulation for each CS with a cursor on a visual analogue scale from 0 = “no expectancy” to 100 = “high expectancy” before the start and after the end of each experimental phase on each day. The start position of the cursor on the scale was determined randomly.

### Acquisition of fMRI data

FMRI data was acquired on a Siemens MAGNETOM Trio 3 Tesla MRI System using a 32-channel head coil. Resting-state and task-based fMRI data were recorded using gradient echo, echo planar imaging (EPI) with a multiband sequence covering the whole brain (TR: 1000 ms, TE: 29 ms, multi-band acceleration factor: 4, voxel-size: 2.5 mm isotropic, flip angle 56°, field of view: 210 mm; (46)). A high-resolution T1 weighted image was acquired after the experiment on day 1 for anatomical visualization and normalization of the EPI data (TR: 1900 ms, TE: 2540 ms, voxel size: 0.8 mm isotropic, flip angle 9°, field of view: 260 mm). T2 weighted images were collected for preventative neuro-radiological diagnostics for all participants (45 slices, TR: 6100 ms, TE: 79 ms, voxel size: 3 mm isotropic, flip angle: 120°). Lastly, we collected multidimensional diffusion-weighted tensor images from each participant (72 slices, voxel-size: 2mm isotropic, TR: 9100 ms, TE: 85 ms, number of directions: 64, b-value: 1000s/mm2).

### Preprocessing of fMRI data

FMRI data was preprocessed and analyzed using statistical parametric mapping (SPM12, Wellcome Trust Centre for Neuroimaging, London, UK, http://www.fil.ion.ucl.ac.uk/) running on Matlab 2017b (MathWorks®, Natick, Massachusetts, USA). The first 5 volumes of each scan were discarded due to equilibrium effects. Preprocessing included realignment and co-registration of the mean functional image to the T1 weighted anatomical image. Subsequently, the T1 weighted anatomical image was segmented and normalized to Montreal Neurological Institute (MNI) space based on SPM’s Unified Segmentation. Normalization of the functional images was achieved by applying the resulting deformation fields to the realigned and co-registered functional images. Lastly, the functional data was smoothed using a 4mm full-width-at-half-maximum Gaussian smoothing kernel. Data of participants was excluded when movement peaks exceeded more than 3 mm or 2°.

### Data exclusion criteria

SCR data was only excluded when participants had zero valid SC responses (n=8), or due to technical problems (n=7). Furthermore, MRI data was excluded due to motion artefacts (n=2), incomplete data (n=13), participant falling asleep (n=1), positive drug test (n=2), and for dataset 4 all participants receiving an L-Dopa pill before extinction (n=30), see Table 1. This led to datasets of n=173 (SCR data) and n=122 (MRI data).

**Table 1.** Participant exclusions. Out of a total of 234 recruited participants, 61 participants were excluded from the LCGM analysis of SCR data (a). 51 participants had no (valid) fMRI data analysis (b).

| Exclusion reasons | Data set 1<br>original<br>N=45 | Data set 2<br>original<br>N=70 | Data set 3<br>original<br>N=56 | Data set 4<br>original<br>N=63 |
| --- | --- | --- | --- | --- |
| **a) Exclusions for LCGM analysis** |  |  |  |  |
| Technical problems SCR data | n=0 | n=2 | n=3 | n=2 |
| Zero valid SCR | n=3 | n=0 | n=1 | n=4 |
| Incomplete data | n=5 | n=2 | n=1 | n=5 |
| Asleep during experiment | n=0 | n=1 | n=0 | n=0 |
| Positive drug test | n=0 | n=0 | n=2 | n=0 |
| L-DOPA before extinction | n=0 | n=0 | n=0 | n=30 |
| **Final subjects for LCGM analysis**<br>n=173 | n=37 | n=65 | n=49 | n=22 |
| **b) Exclusions for fMRI analysis** |  |  |  |  |
| MRI data, motion artefacts | n=0 | n=0 | n=N.A. | n=2 |
| **Final subjects for fMRI analysis**<br>n=122 | n=37 | n=65 | n=0 | n=20 |

### Public Datasets

Datasets from https://osf.io/rkp3a/ were considered for LCGM based on a predefined set of inclusion criteria. SCR data had to be available on a trial-by-trial level, and the dataset was required to contain an extinction phase. Furthermore, only datasets using a differential conditioning design (i.e., with CS+ and CS− stimuli) were considered. Datasets were excluded if the number of CS+ and CS− trials was unequal within a single phase, or if fewer than 10 CS trials were included in any single phase (conditioning and extinction). To ensure consistency in learning trajectories, datasets that involved any intervention before the end of the extinction phase were also excluded. Finally, only datasets with a sample size greater than 100 participants were eligible for analysis. This led to the inclusion of processed datasets of Chalkia et al. (30) and Lonsdorf et al. (31).

### Latent Class Growth Model Analysis

LCGM is a data-driven, explorative statistical method and was applied on SCR data using the *hlme* function in R 3.5.9 for estimating latent class linear mixed models. LCGM analysis characterizes individual differences in parameters, reflecting participants’ changes in outcomes over time. Individuals are classified into latent classes predicated on similar patterns over time (47,48). Single trials per participant on day 1 and day 2 grouped into CS+ and CS-separately for the in-house datasets, Chalkia et al. dataset, and Lonsdorf et al. dataset were included in the models. Day 3 trials were ignored, such as to assure identical classification criteria in all data sets. Furthermore, participants who received L-DOPA before extinction in experiment 4 (n=30) were not included. Reducing the odds of convergence towards a local maximum and optimizing the reliability of the log-likelihood estimation, 30 iterations from 100 vectors of initial values were run. The estimation procedure was then finalized only for the departure that provided the best log-likelihood after 30 iterations. Models were run for one, two, three, and four classes. Model selection was based on these criteria: Apparent drop of Bayesian Information Criterion (BIC) and Akaike Information Criterion (AIC) (49), a large entropy score, class proportion of >5%, and the smallest number of classes that would still be theoretically meaningful. Participants received a probability score for each trajectory and were assigned to a class based on the highest probability score.

### Statistical analysis SCR

To analyze differential learning and extinction, SCRs were averaged per trial type (CS+, CS−), time (first and last 20% of trials), and phase (conditioning, extinction, test) for each participant, following a prespecified procedure systematically applied by our lab in our extinction studies (32–34). Repeated-measures ANOVAs were conducted separately for each phase, with stimulus (CS+, CS−) and time (early, late) as within-subject factors and class (1, 2) as a between-subject factor. Interaction effects (stimulus by time by class) were used to assess differences in learning trajectories between classes. Follow-up tests were conducted within each class where appropriate.

#### Supplementary analyses

For the analyses of our laboratory datasets, we examined whether trait anxiety (STAI-T) or study membership predicted class membership. This was tested using logistic regression with class membership as the dependent variable and STAI-T and study as predictors. As a follow-up, we also ran a repeated-measures ANOVA on SCR of Day 2 including study to assess potential moderation of class-related effects. For the two publicly available SCR datasets, we tested whether sex predicted class membership by fitting a logistic regression model with class membership as the dependent variable and sex as the predictor.

### Single-subject level analysis of fMRI data during extinction

First-level analyses were implemented using a general linear model for day 2 with regressors for CS+ and CS- onsets and offsets in bins of five trials, ratings, and context onsets. All regressors were modelled as delta-functions and convolved with the canonical hemodynamic function (HRF). FAST correction for autocorrelation and a high-pass filter (128 sec) was applied.

### Striatal prediction error activity analysis

Second-level analyses were implemented in SPM12 using a flexible factorial design with the factors time (early, mid, late; within-subject) and class (class 1 vs. class 2; between-subject), with subject modeled as a factor to account for repeated measures. The model tested CS+ vs. CS-offset contrasts from the first level, specifically examining the interaction between class and time. Analyses were conducted at the whole-brain level, with additional small volume correction (SVC), using a mask based on the whole-brain meta-analysis by Garrison et al. (50) and restricted to striatal voxels (caudate, putamen, pallidum, and nucleus accumbens) as implemented and provided by Thiele et al. (15). SVC was applied at peak level. Peaks were only regarded as significant when falling below an initial uncorrected voxel threshold of 0.01 and a family-wise error (FWE)-corrected peak threshold of 0.05 adjusted for the small volume. Based on this analysis, a spherical ROI (6 mm) with the peak voxel [-14 10 -8] was created for extraction of contrast and beta estimates, for the purpose of effect illustration.

### Statistical analysis fMRI

To assess stimulus-specific offset responses during extinction, contrast estimates (CS+ > CS−) and individual beta values were extracted from predefined time windows (early, mid, late extinction). Group-level differences in contrast estimates were evaluated using post-hoc Welch’s t-tests across time points. Additionally, repeated-measures ANOVAs were conducted on beta estimates with stimulus (CS+, CS−) and time (early, mid, late) as within-subject factors, separately for each class. All tests were two-sided.

## Results

### Heterogeneity in extinction learning

To characterize heterogeneity in extinction learning, we reanalyzed a set of four Pavlovian conditioning and extinction experiments conducted in N=234 healthy participants (Fig. 1). Participants were conditioned on day 1 to a visual stimulus (CS+), applying an electric stimulus as a reinforcement (US) at stimulus offset (reinforcement rate of 50%). One day later (day 2), participants returned for an extinction session. Three of the studies also tested for extinction memory retrieval one day after the extinction session (day 3). In all studies, conditioned fear was assessed by SCRs towards the conditioned (CS+) relative to an unreinforced control stimulus (CS-). LCGM analysis of trial-wise SCRs was restricted to the conditioning and extinction data from days 1 and 2, to not confound individual differences in extinction learning with potential differences in extinction memory retrieval, which is a partly independent process (32).

Because experiments had been performed for the purpose of investigating the effect of the drug L-DOPA on extinction learning and memory, all participants were male (32–34). L-DOPA or placebo were administered either before or after extinction on day 2. Participants having received L-DOPA before extinction were excluded.

FMRI data was collected in three studies. Application of exclusion criteria led to a final pooled sample of n=173 for the LCGM analysis and n=122 for the fMRI analysis (see Table 1 and Methods).

The LCGM analysis was run for 1-, 2-, 3-, and 4-class models (Table 2). Model selection was based on the following criteria: A large decrease of Akaike Information Criterion (AIC) and Bayesian Information Criterion (BIC), a large entropy score, single class proportion >5%, and the smallest number of classes that would still be theoretically meaningful (48,49). The 2-class model was chosen for further analysis since AIC and BIC of the 2-class model showed the biggest drop from model 1 and the highest entropy score. Furthermore, it is the lowest number of theoretically meaningful classes. The larger class (class 2) contained 75.7% of participants (n=131) and the smaller class (class 1) 24.3% (n=42). See Fig. 2. Visual inspection of class profiles shows absent extinction of CRs in class 1 and full extinction in class 2. Classes already diverge on day 1, where they appear to show comparable differential conditioning (CS+>CS-differences), but class 2 reaches lower average SCR levels than class 1 as conditioning progresses.

**Figure 2.**
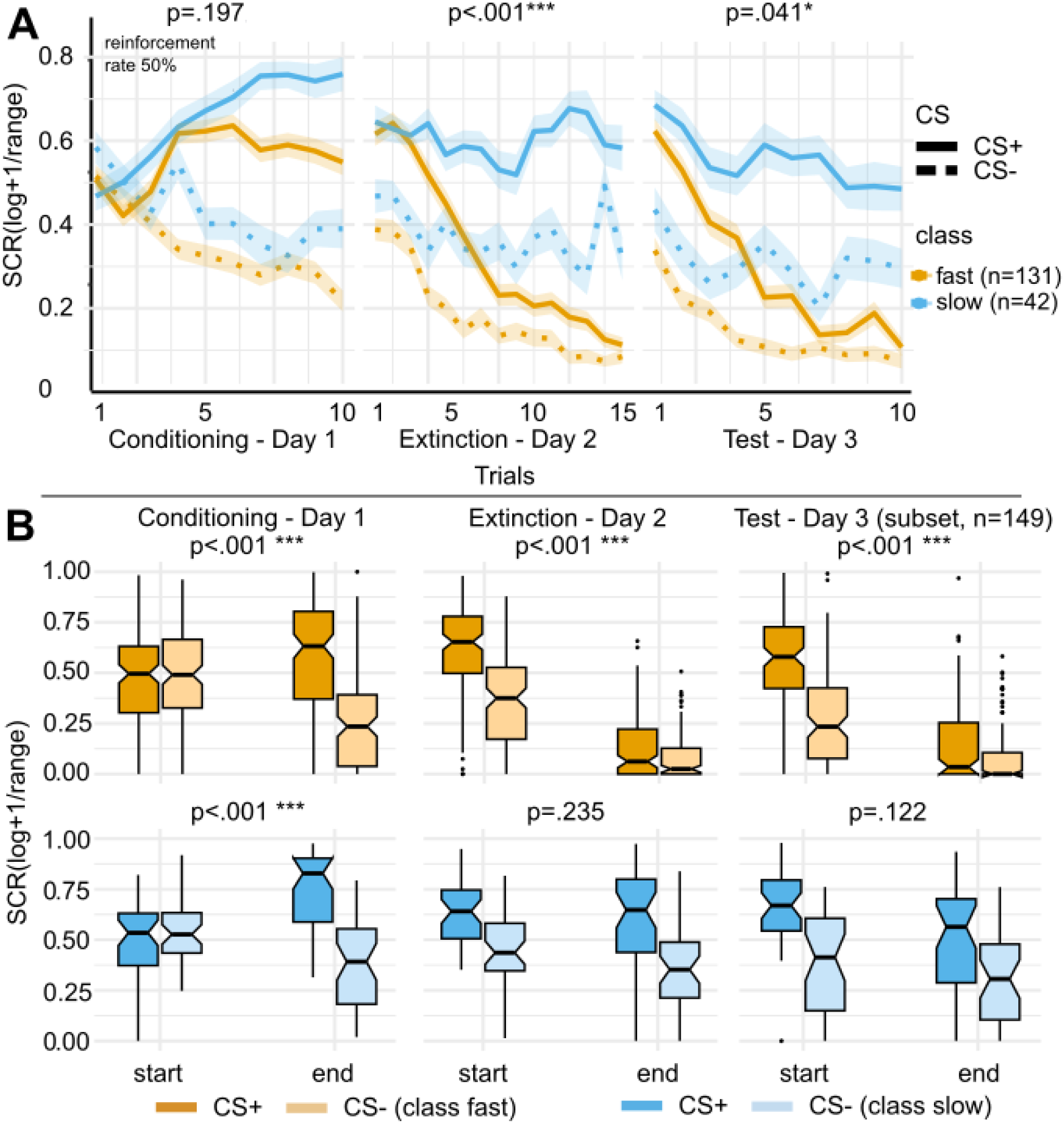
Differential SCR dynamics across conditioning, extinction, and test phases in LCGM-defined extinction phenotypes *(fast/slow extinguishers, n=131/42).* **(A)** Skin conductance responses (SCR) across trials during conditioning (Day 1), extinction (Day 2), and test (Day 3; subset undergoing extinction memory retrieval test consists of n=149 (fast/slow extinguishers n=110/39)), shown separately for fast (yellow, n=131) and slow (blue, n=42) extinguishers identified via LCGM analysis of day 1 and day 2 data. Solid and dashed lines indicate CS+ and CS− responses, respectively. Shaded areas represent ± standard error of the mean (SEM). P-values reflect the three-way ANOVA interaction: stimulus (CS+ vs. CS−) by time (early vs. late) by group (fast vs. slow). **(B)** Boxplots of SCRs at the beginning (start) and end of each phase, shown separately for stimulus type (CS+ dark, CS-light) and group. Top row: fast extinguishers; bottom row: slow extinguishers. P-values indicate ANOVA stimulus by time interaction effects. Data presented as median ± 1.5*IQR.

**Table 2.** LCGM classification of trial-wise SCR time courses from days 1 and 2 (n=173). Classification was based on factors trial number and CS type. loglik = log-likelihood of a latent class model, AIC = Akaike Information Criterion, BIC = Bayesian Information Criterion. Best fitting model values are displayed in bold.

| Number of classes | loglik | AIC | BIC | Entropy | %class1 | %class2 | %class3 | %class4 |
| --- | --- | --- | --- | --- | --- | --- | --- | --- |
| 1 | -1408.35 | 2838.69 | 2873.38 | 1.0 | 100.0 |  |  |  |
| **2** | **-1391.02** | **2812.04** | **2859.34** | **0.80** | **24.28** | **75.72** |  |  |
| 3 | -1384.46 | 2806.91 | 2866.83 | 0.66 | 34.68 | 40.46 | 24.86 |  |
| 4 | -1380.46 | 2806.93 | 2879.45 | 0.66 | 6.94 | 24.28 | 36.99 | 31.79 |

**Table 3.** LCGM classification of trial-wise SCR time courses from days 1 and 2 of the Chalkia dataset (upper table, n=165) and the Lonsdorf dataset (lower table, n=110). Classification was based on factors trial number and CS type. loglik = log-likelihood of a latent class model, AIC = Akaike Information Criterion, BIC = Bayesian Information Criterion. Best fitting model values are displayed in bold. AIC and BIC of the 2-class models showed a drop from model 1 values and the highest entropy score (only Lonsdorf). Class 1 of the 3-class solution in the Chalkia dataset was <5%, therefore also the 2-class model was chosen.

| Number of classes | loglik | AIC | BIC | Entropy | %class1 | %class2 | %class3 |
| --- | --- | --- | --- | --- | --- | --- | --- |
| 1 | 1537.12 | -3050.24 | -3012.97 | 1 | 100 |  |  |
| **2** | **1545.22** | **-3058.44** | **-3008.74** | **0.74** | **84.24** | **15.76** |  |
| 3 | 1550.88 | -3061.77 | -2999.65 | 0.85 | 1.21 | 15.76 | 83.03 |
| 1 | 1683.39 | -3342.78 | -3310.16 | 1.0 | 100.0 |  |  |
| **2** | **1699.31** | **-3366.61** | **-3323.12** | **0.89** | **20.54** | **79.46** |  |
| 3 | 1705.66 | -3371.31 | -3316.94 | 0.83 | 58.93 | 25.89 | 15.18 |

To quantify differences between classes in the development of differential learning during conditioning and extinction, we calculated stimulus (CS+, CS-) by time (20% trials at start of phase and end of phase, see Methods, as prespecified by (32–34)) by group (class 1, class 2) interaction effects for each phase (see full statistics in Supplementary Tables 1-6). See Fig. 2B. Differential learning at conditioning was comparable in both classes (repeated-measures ANOVA: stimulus by time by group interaction: F_1,171_=2.48, p=.12, generalized η^2^=.002, n=173). Differential extinction strongly differed (stimulus by time by group interaction: F_1,171_=18.47, p<.001, generalized η^2^=.02, n=173), with no stimulus by time effect in class 1 (F_1,41_=1.45, p=.24, generalized η^2^=.007, n=42). Data from the subset of participants that had undergone an extinction memory test on day 3 (see Fig. 1: data sets 1-3; n=149) also showed differential effects, with stronger decline of CRs in class 2 (stimulus by time by group interaction: F_1,147_=4.16, p=.04, generalized η^2^=.004). In class 1, although the stimulus by time interaction was not significant (F_1,38_= 2.50, p=.12, generalized η^2^=.01, n=39), there was a clear main effect of stimulus (F_1,38_= 54.51, p<.001, generalized η^2^=.19) and time (F_1,38_= 9.58, p=.004, generalized η^2^=.06), indicating an overall decrease in responding. This suggests class 1 was not entirely unable to extinguish their CRs but slower. On the basis of these results, we label class 1 ‘slow extinguishers’ and class 2 ‘fast extinguishers’.

Trait anxiety (STAI-T) did not influence class membership (see statistics in Supplementary Table 7), while class membership showed modest study-related variability (omnibus χ²(3)=13.83, p=.003), with only limited pairwise differences after multiplicity correction (see Supplementary Table 7, see Supplementary Fig. 1). Study membership however had no main effect on SCR and did not moderate the critical interaction effect on Day 2 (four-way p=.727, see Supplementary Table 8), indicating the key class-related pattern was robust across studies.

To assess the replicability of learning types derived from trial-wise SCR data across two-day differential conditioning and extinction protocols, we applied the same LCGM procedure to two publicly available datasets with at least n>100 female and male healthy participants (30,31); see Methods). Although the datasets differed in reinforcement rates during conditioning (37.5% in Chalkia et al., 100% in Lonsdorf et al.), a 2-class solution was again selected in both based on the same model selection criteria. Visual inspection of class profiles revealed consistent patterns: The slow extinguisher classes (class 2 Chalkia et al., class 1 Lonsdorf et al.) showed reduced extinction of CRs, while the fast extinguisher classes (class 1 Chalkia dataset, class 2 Lonsdorf dataset) exhibited full extinction (see full statistics in Supplementary Tables 9-14; 16-21). On day 1, both groups diverged similarly across datasets, showing comparable CS+ > CS− differences (Fig. 3), although fast extinguishers reached lower average SCR levels than slow extinguishers as conditioning progressed. Importantly, the Chalkia dataset showed a significant three-way interaction (stimulus by time type by group: F_1,163_=9.46, p=.002), confirming that extinction trajectories differed by class. In contrast, the Lonsdorf dataset showed only a non-significant trend-level interaction (F_1,108_=3.17, p=.078), possibly because the 100% reinforcement during conditioning made US omission during extinction more salient, leading to faster and more uniform extinction and thus reduced between-subject variability. In both datasets, sex did not influence class membership (see full statistics in Supplementary Tables 15 and 22).

**Figure 3.**
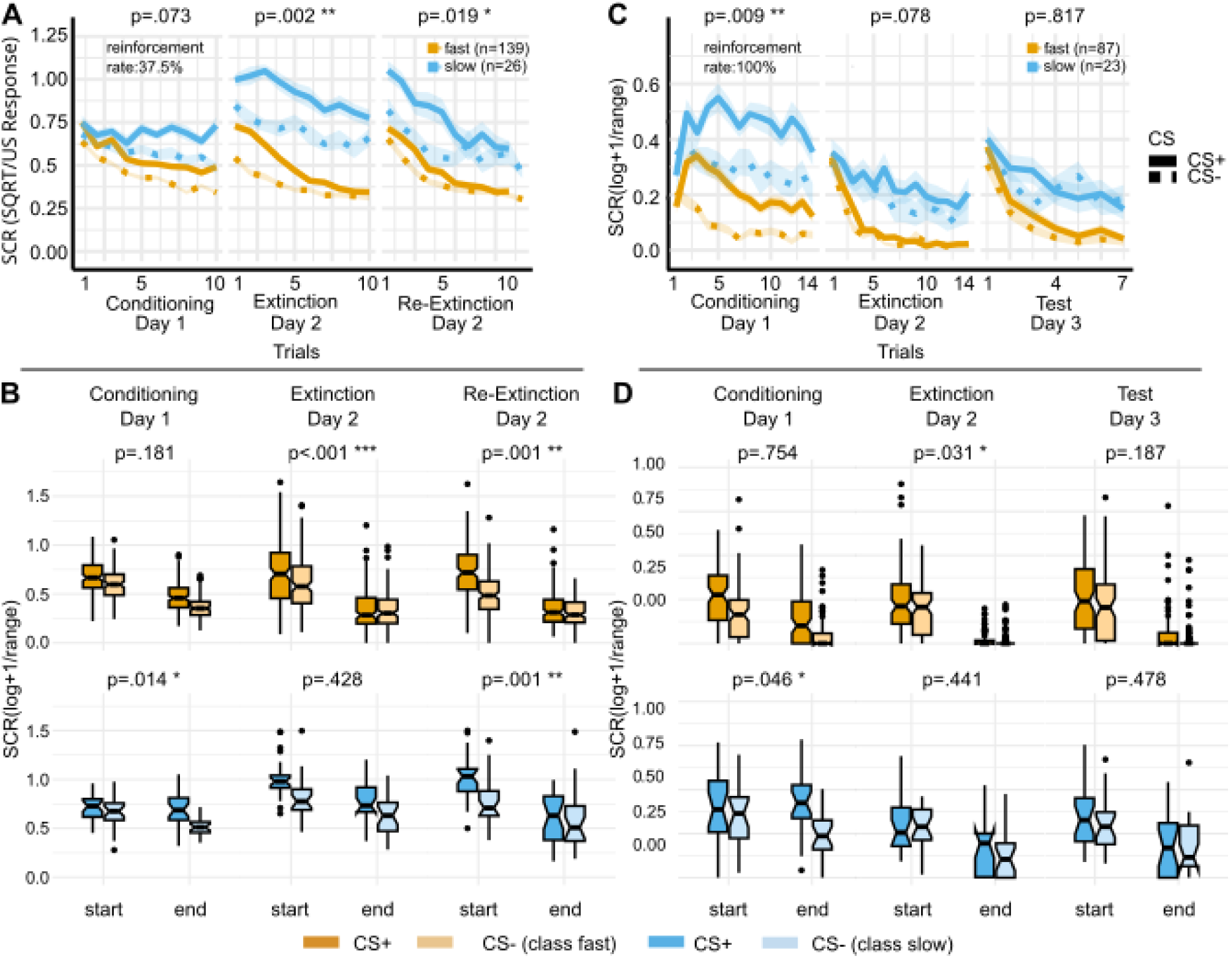
Differential SCR dynamics across conditioning, extinction, and test phases in LCGM-defined extinction phenotypes in the Chalkia and Lonsdorf datasets. **(A/C)** Skin conductance responses (SCR) across trials during conditioning, extinction, and re-extinction / test, shown separately for fast (yellow) and slow (blue) extinguishers identified via LCGM (A Chalkia dataset, C Lonsdorf dataset). Solid and dashed lines indicate CS+ and CS− responses, respectively. Shaded areas represent ± standard error of the mean (SEM). P-values reflect the three-way ANOVA interaction: stimulus (CS+ vs. CS−) by time (early vs. late) by group (slow vs. fast). **(B/D)** Boxplots of SCRs at the beginning (start) and end of each phase, shown separately for stimulus type (CS+ dark, CS- light) and group. Top row: fast extinguishers; bottom row: slow extinguishers (B Chalkia dataset, D Lonsdorf dataset). P-values indicate ANOVA stimulus by time interaction effects. Data presented as median ± 1.5*IQR.

On the basis of these results, we conjecture that individual differences in fear extinction in healthy humans are well captured by a normative fast extinguishing and a less frequent slow extinguishing type.

### Individual differences in ventral striatal fMRI responses to US omission in extinction

Next, we hypothesized that distinct extinction learning types would also show distinct neural correlates. As extinction learning is thought to be initiated and driven by PE signals generated in the VS by unexpected US omission (12,13,15), the subset of participants with fMRI data (n=122, fast/slow extinguishers: n=98/24; datasets 1,2, and 4; see Table 1 for exclusions, Fig. 4A for SCR time courses), was tested for class differences in CS offset responses in a pre-defined VS region of interest (ROI). The ROI included brain areas identified in a meta-analysis of RPEs (50) but was restricted to striatal voxels by overlapping with the Harvard-Oxford masks for caudate, putamen, pallidum, and nucleus accumbens. It was previously successfully employed by us to identify EPEs in a computational fMRI study using the Rescorla-Wagner model (15). To reduce noise, extinction trials were averaged in five-trial bins (early, mid, late extinction). This analysis revealed a significant stimulus (CS+ vs. CS−) by time (early vs. late extinction) by group (fast vs. slow extinguishers) interaction in a peak in the VS (left pallidum/ventral putamen) at MNI coordinates x,y,z=-14,10,-8 (*Z*=5.21, k_E_=37, *p_SVC_*<.0001 after small-volume correction for multiple comparisons, Fig. 4B).

**Figure 4.**
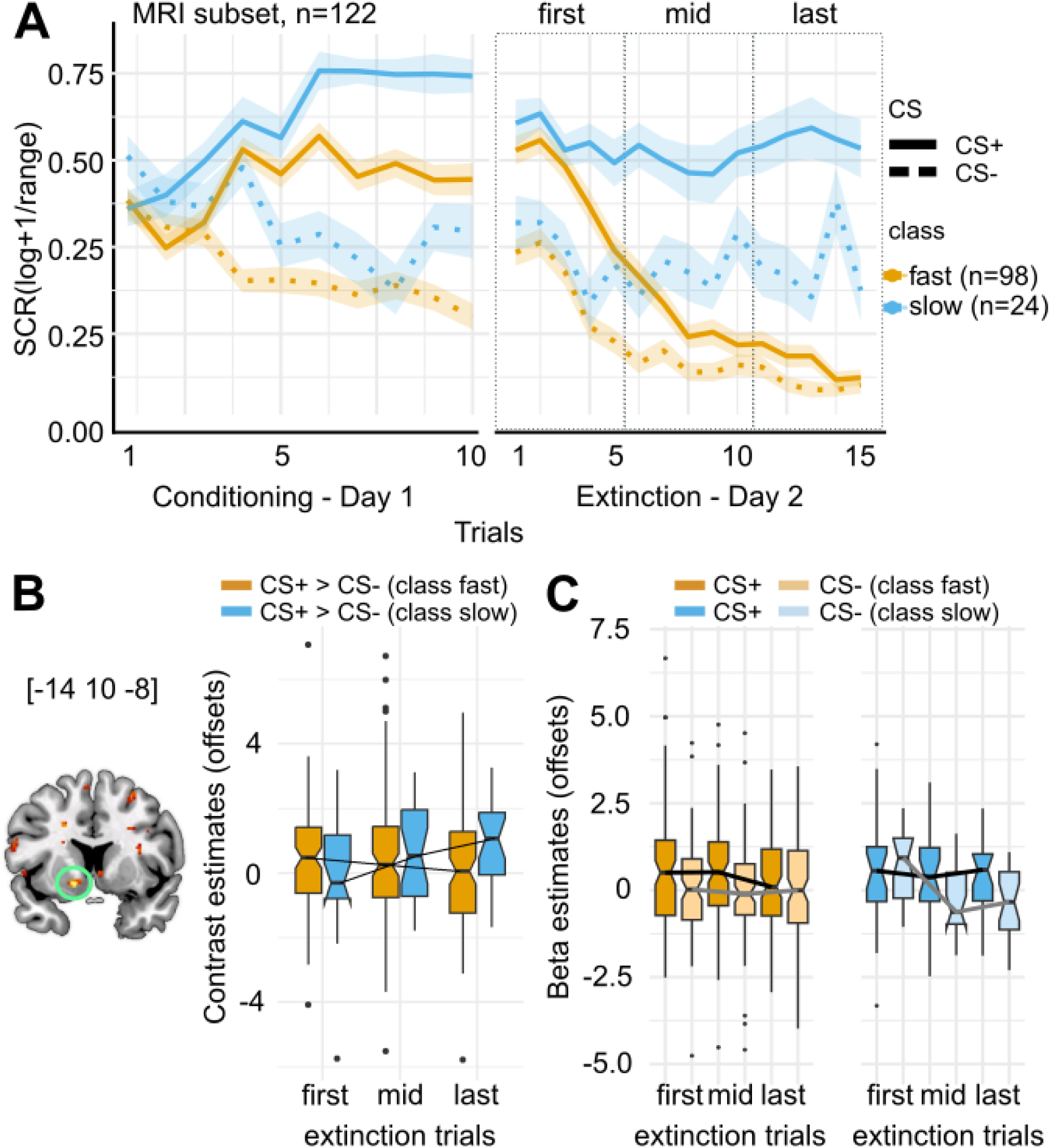
Activity in a striatal mask was investigated in a subgroup of participants undergoing the paradigm in fMRI. **A.** Conditioned responses as SC responses to the CS+ and CS- during conditioning and extinction of a subset used for further MRI analysis (n=122, fast/slow extinguishers, n=98/24). Data is presented as mean ± standard error of the mean (sem). **B.** CS+>CS- offset contrast identified a peak in the ventral striatum (left panel). Contrast estimates show a significant interaction (middle panel). **C.** Separate CS+ (dark) and CS- (light) beta estimates are depicted in the right panel for fast (yellow) and slow (blue) extinguishers. Data presented as median ± 1.5*IQR.

The same peak was also significant in an exploratory whole-brain analysis corrected at p<0.05 (Z=5.21, k_E_=117, p_fwe_=.014). The peak was located in the same position as in the earlier computational EPE analysis by Thiele et al. (15), where it had also been significant both at the ROI and the whole-brain level. The CS+>CS-contrast estimates in Fig. 4B suggest an early differential CS offset signal in the fast extinguishers (class 2), which declines over the course of extinction, conformal with an EPE. A comparable signal is only observed later in the slow extinguishers (class 1). Post-hoc t-tests revealed a significant difference in late extinction (early: *t*_34.18_=1.21, *p*=.24; mid: *t*_44.34_=0.55, *p*=.59; late: *t*_46.57_=2.10, *p*=.04; n=122, Fig. 4B). Whereas fast extinguishers exhibited a stimulus main effect (post-hoc ANOVA: stimulus: *F*_1,97_=8.34, *p*=.005, generalized η^2^=0.02; time: *F*_2,194_=0.55, *p*=.58, generalized η^2^=0.002; stimulus by time: *F*_2,194_=0.70, *p*=.50, generalized η^2^=0.002; n=98), slow extinguishers showed a significant main effect of time and a stimulus by time interaction effect (stimulus: *F*_1,23_=3.87, *p*=.06, generalized η^2^=0.03; time: *F*_2,46_=6.24, *p*=.004, generalized η^2^=0.07; stimulus by time: *F*_2,46_=3.79, *p*=.03, generalized η^2^=0.04; n=24).

## Discussion

In this study, we applied LCGM to trial-wise SCRs during fear conditioning and extinction in four datasets collected in our lab and two publicly available samples. Across all datasets, we consistently identified two groups differing in the time course of CR decline during extinction. One group showed a rapid reduction in differential SCRs (“fast extinguishers”), while the other group exhibited sustained responses and only delayed reduction (“slow extinguishers”). Importantly, we found that these behavioral phenotypes were reflected in ventral striatal activity measured with fMRI: fast extinguishers showed an early CS+ > CS− offset signal during extinction, while slow extinguishers displayed a comparable differential signal only later. These results suggest that individual differences in extinction learning relate to differences in the timing of a ventral striatal signal consistent with an EPE.

This early offset-related CS+ > CS− activation observed in the VS of fast extinguishers aligns with the theoretical profile of an RPE–like signal during extinction learning (10,15). Since omission of an expected aversive outcome constitutes a better-than-expected event, this signal has been interpreted as an appetitive or relief-like PE. That this signal peaked early supports its functional identity as an EPE. By contrast, the delayed emergence of this signal in slow extinguishers suggests a less efficient detection or utilization of US omission early in extinction. One interpretation is that dopaminergic PE signaling is attenuated or delayed in this group, possibly requiring a higher threshold of expectancy violation or a longer accumulation of evidence before a reliable EPE is generated. Alternatively, the late-emerging ventral striatal signal in slow extinguishers could reflect a compensatory or downstream process rather than a canonical EPE. For example, other brain regions such as the ventromedial prefrontal cortex (vmPFC) might initially detect the absence of threat through top-down mechanisms, subsequently activating the VS indirectly once safety information is established (32,33).

Our use of LCGM to characterize extinction trajectories adds an important methodological strength to these findings. The approach allowed us to identify robust subgroups based purely on SCR trajectories, without a priori assumptions about learning models or response shapes. This data-driven strategy is particularly advantageous in extinction research, where trial-by-trial dynamics may deviate from the strictly monotonic curves assumed in associative frameworks. By not enforcing a single learning function, LCGM captures heterogeneity that may arise from thresholded updating, shifts in contingency awareness, or contextual influences.

Nevertheless, LCGM also comes with limitations. It describes surface-level behavioral patterns rather than inferring latent cognitive mechanisms. For instance, differences between fast and slow extinguishers may stem from explicit cognitive inference (e.g., belief updating about contingencies) or more automatic processes (e.g., stimulus salience or arousal), which LCGM alone cannot disambiguate. Moreover, class assignment can be sensitive to minor differences in trajectory shapes. Importantly, classification outcomes may also vary with reinforcement schedule: while our datasets used partial (50%) reinforcement, the Chalkia sample used 37.5% and the Lonsdorf dataset 100%, and the critical stimulus by time by group interaction was only trend-significant in the 100% condition, likely because learning is more straightforward under continuous reinforcement.

The overall validity of our 2-class solution is further supported by prior work. Gazendam et al. (27), using a larger sample, identified similar fast and slow extinction profiles – two groups with rapid extinction and low overall SCRs, closely resembling our fast extinguishers, and one group with high SCR reactivity and poor extinction, paralleling our slow extinguishers. Notably, their study also revealed an additional “low-responder” class, characterized by minimal SCR reactivity throughout, which was not present in our data. These findings suggest that richer class structures may emerge in larger or more heterogeneous samples, possibly capturing more nuanced extinction phenotypes.

Relatedly, some individuals may show globally elevated SCRs across phases of the experiment, pointing to a hyper-reactive autonomic profile. This could reflect high trait arousal that interferes with extinction. Supporting this notion, Andres et al. (33) found that elevated baseline salivary alpha-amylase (sAA) – a proxy for basal sympathetic nervous system activity – was associated with impaired extinction learning and memory retrieval. In our context, the slow extinguisher class may reflect such a high-arousal phenotype, possibly driven by noradrenergic tone interfering with vmPFC function or striatal signaling. Given that LCGM captures both SCR amplitude and trajectory, tonic arousal could bias class membership independently of learning-specific processes. Future work could address this by incorporating trait-level physiological markers, such as sAA or resting heart rate variability, which could practically be used clinically for pre-treatment diagnostics and stratification of therapy.

Another important consideration is the role of sex. Because our lab-based datasets included only male participants due to pharmacological constraints (see Methods), the generalizability of our findings is limited. However, similar class structures in the mixed-sex Chalkia, Lonsdorf and Gazendam datasets (27,30,31) suggest that the observed extinction phenotypes are not restricted to males. Nonetheless, future studies should directly test for sex differences in both behavioral and neural indicators of extinction learning.

Beyond theoretical implications, our findings also have clear clinical relevance. Extinction learning is the foundational mechanism behind exposure-based therapies for the prevention (51) and treatment (52) of anxiety disorders. Our slow extinguisher phenotype, marked by delayed ventral striatal responses to US omission, may index individuals at greater risk for poor treatment outcomes. This notion is consistent with clinical findings showing that delayed or incomplete extinction predicts higher relapse and lower treatment efficacy (53–55).

Accordingly, our results offer two translational pathways. First, SCR-based subgrouping, or trait-level physiological indices, such as elevated sAA, could be used to stratify patients before treatment, identifying those who might benefit from tailored interventions such as pharmacological augmentation or extended therapy sessions. Second, the ventral striatal blood oxygen level dependent (BOLD) response to CS+ offset may serve as a candidate neural biomarker for exposure therapy success. While SCRs are logistically easier to implement, their test–retest reliability is limited. As demonstrated by Klingelhoefer-Jens et al. (56), SCR patterns do not replicate well within individuals over time, whereas fMRI patterns during acquisition are more stable. This suggests that BOLD signals – despite being resource-intensive – may offer more reliable individual-level markers for therapeutic monitoring.

Future research should also explicitly examine how neuromodulatory interventions, such as transcranial ultrasound stimulation (TUS), influence extinction trajectories and neural signatures, potentially enhancing or accelerating ventral striatal responses or modulating network connectivity to promote faster extinction. Incorporating diverse data types such as pupil responses, BOLD activity, and subjective ratings alongside SCRs will further improve characterization and prediction of extinction phenotypes and clinical outcomes.

In conclusion, we identified fast and slow extinguishers based on trial-wise SCR data and showed that these groups differ in the timing of a ventral striatal response to US omission during extinction. These findings support the role of EPE-like signaling in driving CR reduction and highlight the value of combining data-driven behavioral clustering with fMRI to uncover neurocomputational mechanisms of individual learning differences. Such approaches may ultimately inform personalized treatments for anxiety and stress-related disorders.

## Acknowledgements

For the published datasets (1–3), we thank B. Meyer, K. Yuen, M. Ilhan-Bayrakcı, A. Schick, J. Behr, P. Seifert, N. Schappe, A. Droby and J. Meier for support with data collection, analysis and blinding. For Dataset 4, we thank M. Thiele, M. Lückel and K. Yuen for substantial contributions to data acquisition and conceptualization.

R.K. Received funding from the Deutsche Forschungsgemeinschaft (DFG; CRC 1193, subproject C01, and subproject B01), the State of Rhineland-Palatinate (DRZ program).

## Author contributions

Experimental Design: E.A., C.-P.H., A.M.V.G., O.T., R.K.; Data acquisition: E.A., C.-P.H., A.M.V.G., O.T.; Conceptualization: E.A. and R.K.; Analysis: E.A.; Visualization: E.A.; Writing – original draft: E.A. and R.K.; Writing review & editing: E.A. and R.K.

## Data availability

Derivative data (trial-by-trial SCR, contrast and beta estimates) will be made available upon publication.

## Supplementary Information

### Andres et al. dataset

**Supplementary Table 1:** Day 1 ANOVA.

| *Effect* | *DFn* | *DFd* | *F* | *p* | *p<.05* | *ges* |
| --- | --- | --- | --- | --- | --- | --- |
| *Class* | 1 | 171 | 13.31 | 3.49e-04 | * | 0.031 |
| *Stimulus* | 1 | 171 | 93.80 | 5.84e-18 | * | 0.06 |
| *Time* | 1 | 171 | 0.09 | 0.75 |  | 0.0001 |
| *Class:Stimulus* | 1 | 171 | 0.04 | 0.83 |  | 0.00003 |
| *Class:Time* | 1 | 171 | 7.28 | 0.007 | * | 0.012 |
| *Stimulus:Time* | 1 | 171 | 123.69 | 5.75e-22 | * | 0.10 |
| *Class:Stimulus:Time* | 1 | 171 | 1.67 | 0.19 |  | 0.001 |

**Supplementary Table 2:**
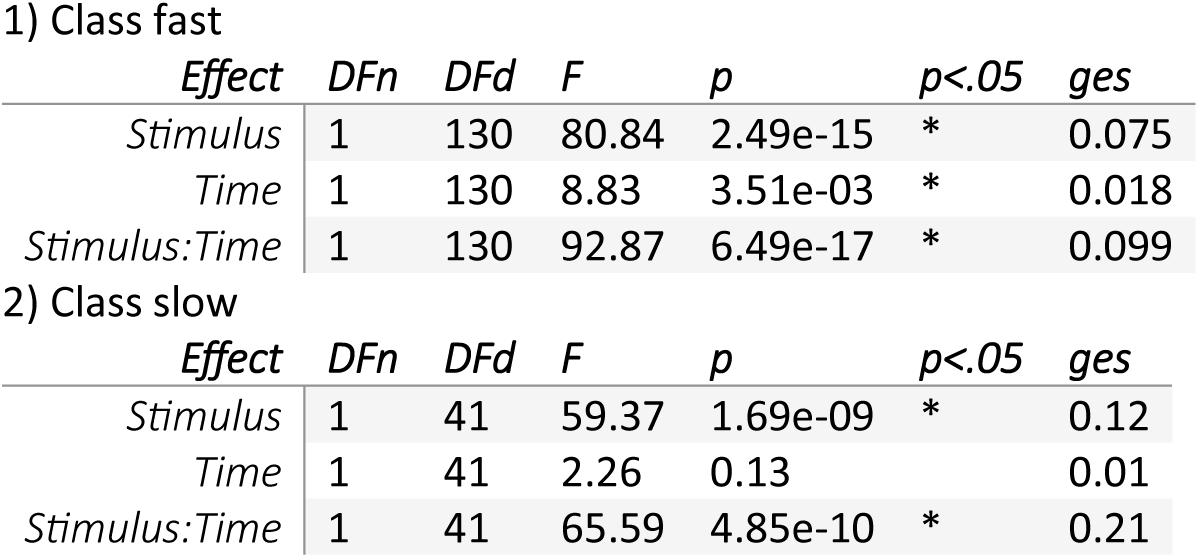
Day 1 Within-Class ANOVAs.

1) Class fast
| *Effect* | *DFn* | *DFd* | *F* | *p* | *p<.05* | *ges* |
| --- | --- | --- | --- | --- | --- | --- |
| *Stimulus* | 1 | 130 | 80.84 | 2.49e-15 | * | 0.075 |
| *Time* | 1 | 130 | 8.83 | 3.51e-03 | * | 0.018 |
| *Stimulus:Time* | 1 | 130 | 92.87 | 6.49e-17 | * | 0.099 |

**Supplementary Table 2: Day 1 Within-Class ANOVAs.**
| *Effect* | *DFn* | *DFd* | *F* | *p* | *p<.05* | *ges* |
| --- | --- | --- | --- | --- | --- | --- |
| *Stimulus* | 1 | 41 | 59.37 | 1.69e-09 | * | 0.12 |
| *Time* | 1 | 41 | 2.26 | 0.13 |  | 0.01 |
| *Stimulus:Time* | 1 | 41 | 65.59 | 4.85e-10 | * | 0.21 |

**Supplementary Table 3:** Day 2 ANOVA.

| *Effect* | *DFn* | *DFd* | *F* | *p* | *p<.05* | *ges* |
| --- | --- | --- | --- | --- | --- | --- |
| *Class* | 1 | 171 | 116.91 | 4.27e-21 | * | 0.18 |
| *Stimulus* | 1 | 171 | 120.39 | 1.51e-21 | * | 0.14 |
| *Time* | 1 | 171 | 167.07 | 4.22e-27 | * | 0.19 |
| *Class:Stimulus* | 1 | 171 | 3.95 | 0.048 | * | 0.005 |
| *Class:Time* | 1 | 171 | 98.64 | 1.22e-18 | * | 0.12 |
| *Stimulus:Time* | 1 | 171 | 3.95 | 0.04 | * | 0.004 |
| *Class:Stimulus:Time* | 1 | 171 | 18.47 | 2.88e-05 | * | 0.02 |

**Supplementary Table 4:**
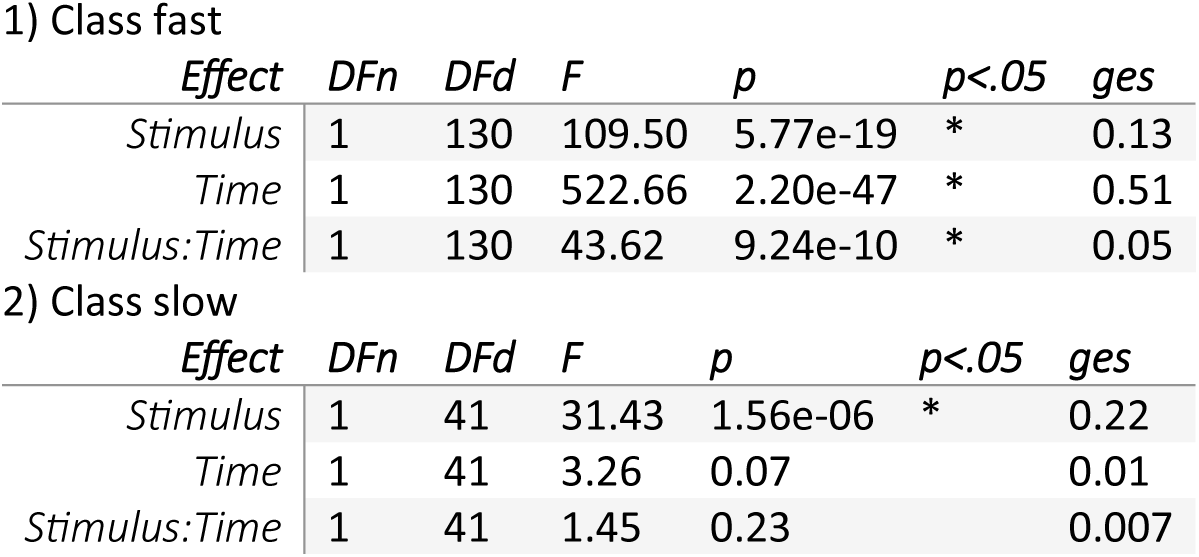
Day 2 Within-Class ANOVA.

1) Class fast
| *Effect* | *DFn* | *DFd* | *F* | *p* | *p<.05* | *ges* |
| --- | --- | --- | --- | --- | --- | --- |
| *Stimulus* | 1 | 130 | 109.50 | 5.77e-19 | * | 0.13 |
| *Time* | 1 | 130 | 522.66 | 2.20e-47 | * | 0.51 |
| *Stimulus:Time* | 1 | 130 | 43.62 | 9.24e-10 | * | 0.05 |

**Supplementary Table 4: Day 2 Within-Class ANOVA**
| *Effect* | *DFn* | *DFd* | *F* | *p* | *p<.05* | *ges* |
| --- | --- | --- | --- | --- | --- | --- |
| *Stimulus* | 1 | 41 | 31.43 | 1.56e-06 | * | 0.22 |
| *Time* | 1 | 41 | 3.26 | 0.07 |  | 0.01 |
| *Stimulus:Time* | 1 | 41 | 1.45 | 0.23 |  | 0.007 |

**Supplementary Table 5:** Day 3 ANOVA.

| *Effect* | *DFn* | *DFd* | *F* | *p* | *p<.05* | *ges* |
| --- | --- | --- | --- | --- | --- | --- |
| *Class* | 1 | 147 | 54.58 | 1.03e-11 | * | 0.12 |
| *Stimulus* | 1 | 147 | 122.00 | 4.97e-21 | * | 0.14 |
| *Time* | 1 | 147 | 117.43 | 1.76e-20 | * | 0.16 |
| *Class:Stimulus* | 1 | 147 | 1.76 | 0.18 |  | 0.002 |
| *Class:Time* | 1 | 147 | 21.28 | 8.53e-06 | * | 0.03 |
| *Stimulus:Time* | 1 | 147 | 23.03 | 3.86e-06 | * | 0.02 |
| *Class:Stimulus:Time* | 1 | 147 | 4.23 | 0.04 | * | 0.004 |

**Supplementary Table 6:** Day 3 Within-Class ANOVA.

| *Effect* | *DFn* | *DFd* | *F* | *p* | *p<.05* | *ges* |
| --- | --- | --- | --- | --- | --- | --- |
| *Stimulus* | 1 | 109 | 88.65 | 9.22e-16 | * | 0.15 |
| *Time* | 1 | 109 | 261.72 | 9.61e-31 | * | 0.35 |
| *Stimulus:Time* | 1 | 109 | 45.19 | 8.49e-10 | * | 0.07 |

**Supplementary Table 6: Day 3 Within-Class ANOVA**
| *Effect* | *DFn* | *DFd* | *F* | *p* | *p<.05* | *ges* |
| --- | --- | --- | --- | --- | --- | --- |
| *Stimulus* | 1 | 38 | 54.51 | 7.49e-09 | * | 0.18 |
| *Time* | 1 | 38 | 9.57 | 0.003 | * | 0.06 |
| *Stimulus:Time* | 1 | 38 | 2.50 | 0.12 |  | 0.009 |

**Supplementary Table 7:**
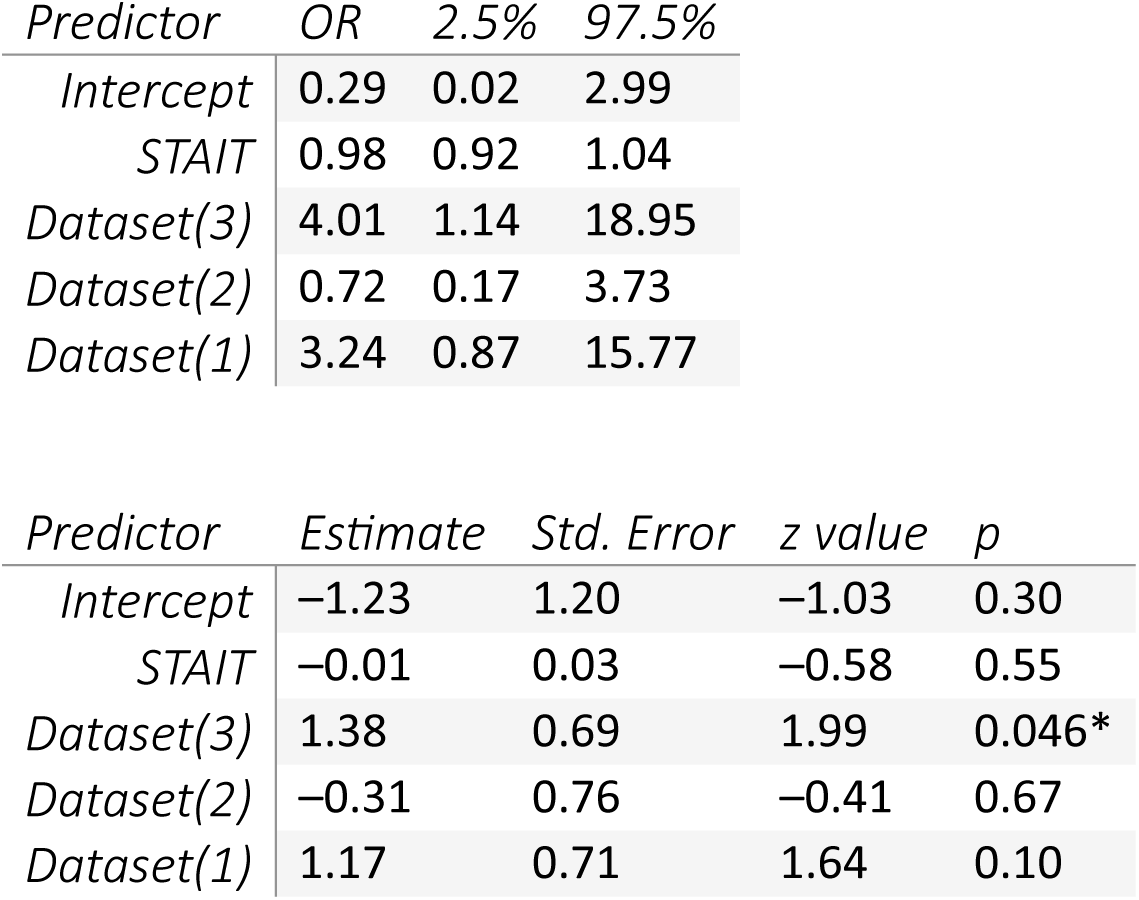
Logistic regression results with class membership as the dependent variable and STAI-T and study as predictors.

| *Predictor* | *OR* | *2.5%* | *97.5%* |
| --- | --- | --- | --- |
| *Intercept* | 0.29 | 0.02 | 2.99 |
| *STAIT* | 0.98 | 0.92 | 1.04 |
| *Dataset(3)* | 4.01 | 1.14 | 18.95 |
| *Dataset(2)* | 0.72 | 0.17 | 3.73 |
| *Dataset(1)* | 3.24 | 0.87 | 15.77 |

| *Predictor* | *Estimate* | *Std. Error* | *z value* | *p* |
| --- | --- | --- | --- | --- |
| *Intercept* | -1.23 | 1.20 | -1.03 | 0.30 |
| *STAIT* | -0.01 | 0.03 | -0.58 | 0.55 |
| *Dataset(3)* | 1.38 | 0.69 | 1.99 | 0.046* |
| *Dataset(2)* | -0.31 | 0.76 | -0.41 | 0.67 |
| *Dataset(1)* | 1.17 | 0.71 | 1.64 | 0.10 |

**Supplementary Figure 1.**
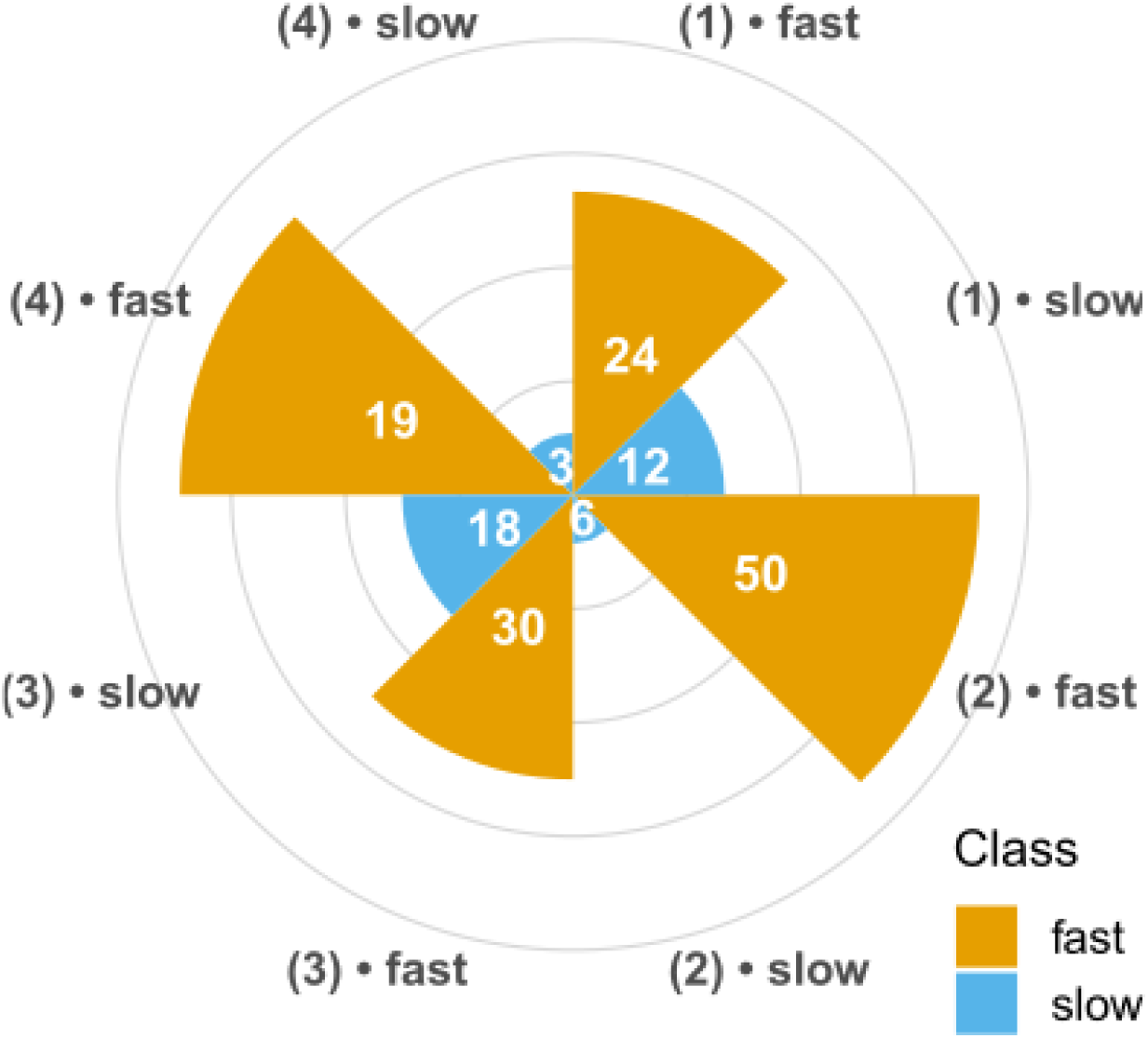
Class membership proportions by study. Rings at 0%, 25%, 50%, 75%, 100% (numbers show n). Datasets 1-4 are as described in Methods and Fig. 1.

**Supplementary Table 8:** Follow-up repeated-measures ANOVA on SCR of Day 2 including study to assess potential moderation of class-related effects.

| Effect | Sum<br>Sq | num<br>Df | Erro<br>r SS | den<br>Df | F value | p-value | Sig |
| --- | --- | --- | --- | --- | --- | --- | --- |
| (Intercept) | 55.74 | 1 | 7.06 | 154 | 1214.93 | < 2.2e-16 | *** |
| Class | 3.66 | 1 | 7.06 | 154 | 79.86 | 1.161e-15 | *** |
| Study | 0.02 | 3 | 7.06 | 154 | 0.15 | 0.92 |  |
| Class:Study | 0.30 | 3 | 7.06 | 154 | 2.22 | 0.08 | . |
| Stimulus | 2.55 | 1 | 5.35 | 154 | 73.62 | 9.540e-15 | *** |
| Class:Stimulus | 0.02 | 1 | 5.35 | 154 | 0.83 | 0.36 |  |
| Study:Stimulus | 0.07 | 3 | 5.35 | 154 | 0.73 | 0.53 |  |
| Class:Study:Stimulus | 0.33 | 3 | 5.35 | 154 | 3.23 | 0.02 | * |
| Time | 3.60 | 1 | 5.53 | 154 | 100.31 | < 2.2e-16 | *** |
| Class:Time | 2.72 | 1 | 5.53 | 154 | 75.93 | 4.352e-15 | *** |
| Study:Time | 0.18 | 3 | 5.53 | 154 | 1.71 | 0.16 |  |
| Class:Study:Time | 0.15 | 3 | 5.53 | 154 | 1.42 | 0.23 |  |
| Stimulus:Time | 0.25 | 1 | 4.34 | 154 | 9.16 | 0.002 | ** |
| Class:Stimulus:Time | 0.14 | 1 | 4.34 | 154 | 5.16 | 0.02 | * |
| Study:Stimulus:Time | 0.29 | 3 | 4.34 | 154 | 3.48 | 0.01 | * |
| Class:Study:Stimulus:Time | 0.03 | 3 | 4.34 | 154 | 0.43 | 0.72 |  |

### Chalkia et al. dataset

**Supplementary Table 9:**
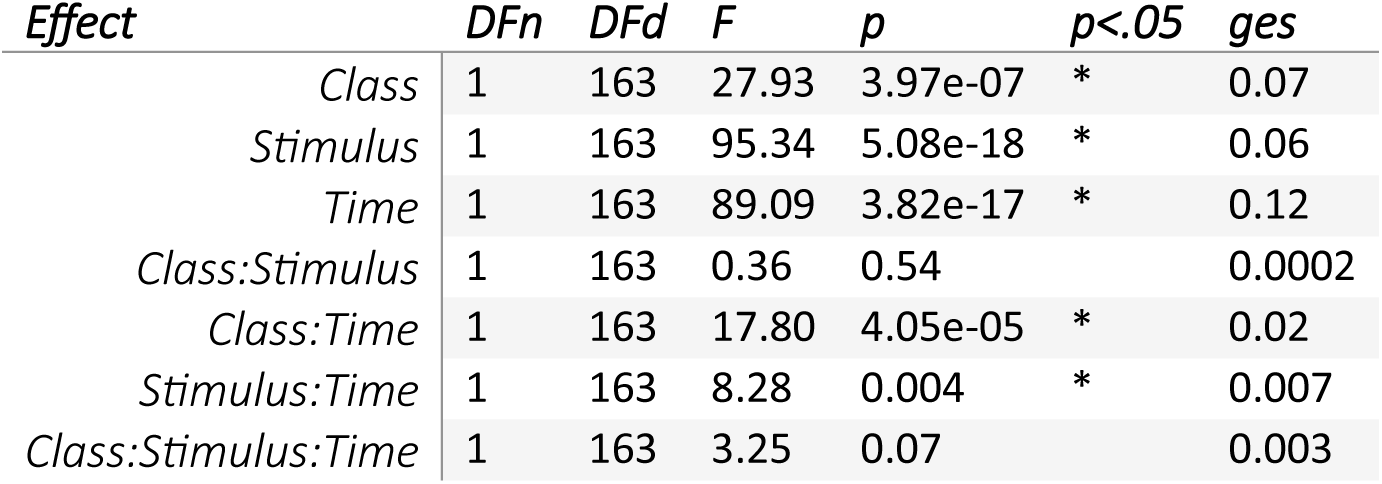
Day 1 ANOVA.

| *Effect* | *DFn* | *DFd* | *F* | *p* | *p<.05* | *ges* |
| --- | --- | --- | --- | --- | --- | --- |
| *Class* | 1 | 163 | 27.93 | 3.97e-07 | * | 0.07 |
| *Stimulus* | 1 | 163 | 95.34 | 5.08e-18 | * | 0.06 |
| *Time* | 1 | 163 | 89.09 | 3.82e-17 | * | 0.12 |
| *Class:Stimulus* | 1 | 163 | 0.36 | 0.54 |  | 0.0002 |
| *Class:Time* | 1 | 163 | 17.80 | 4.05e-05 | * | 0.02 |
| *Stimulus:Time* | 1 | 163 | 8.28 | 0.004 | * | 0.007 |
| *Class:Stimulus:Time* | 1 | 163 | 3.25 | 0.07 |  | 0.003 |

**Supplementary Table 10:**
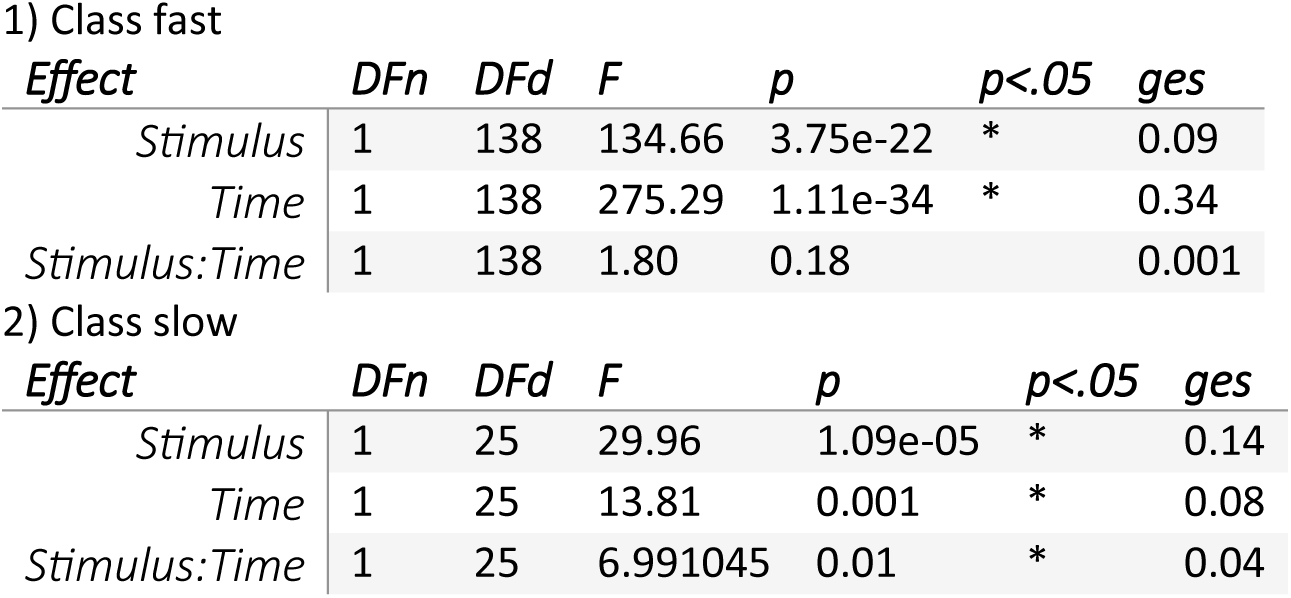
Day 1 Within-Class ANOVA.

1) Class fast
| *Effect* | *DFn* | *DFd* | *F* | *p* | *p<.05* | *ges* |
| --- | --- | --- | --- | --- | --- | --- |
| *Stimulus* | 1 | 138 | 134.66 | 3.75e-22 | * | 0.09 |
| *Time* | 1 | 138 | 275.29 | 1.11e-34 | * | 0.34 |
| *Stimulus:Time* | 1 | 138 | 1.80 | 0.18 |  | 0.001 |

Supplementary Table 10: Day 1 Within-Class ANOVA
| *Effect* | *DFn* | *DFd* | *F* | *p* | *p<.05* | *ges* |
| --- | --- | --- | --- | --- | --- | --- |
| *Stimulus* | 1 | 25 | 29.96 | 1.09e-05 | * | 0.14 |
| *Time* | 1 | 25 | 13.81 | 0.001 | * | 0.08 |
| *Stimulus:Time* | 1 | 25 | 6.991045 | 0.01 | * | 0.04 |

**Supplementary Table 11:** Day 2 ANOVA.

| *Effect* | *DFn* | *DFd* | *F* | *p* | *p<.05* | *ges* |
| --- | --- | --- | --- | --- | --- | --- |
| *Class* | 1 | 163 | 88.32 | 4.92e-17 | * | 0.26 |
| *Stimulus* | 1 | 163 | 109.38 | 6.53e-20 | * | 0.06 |
| *Time* | 1 | 163 | 166.62 | 1.03e-26 | * | 0.14 |
| *Class:Stimulus* | 1 | 163 | 4.24 | 4.08e-02 | * | 0.002 |
| *Class:Time* | 1 | 163 | 4.37 | 3.80e-02 | * | 0.004 |
| *Stimulus:Time* | 1 | 163 | 21.18 | 8.35e-06 | * | 0.008 |
| *Class:Stimulus:Time* | 1 | 163 | 9.45 | 2.46e-03 | * | 0.004 |

**Supplementary Table 12:**
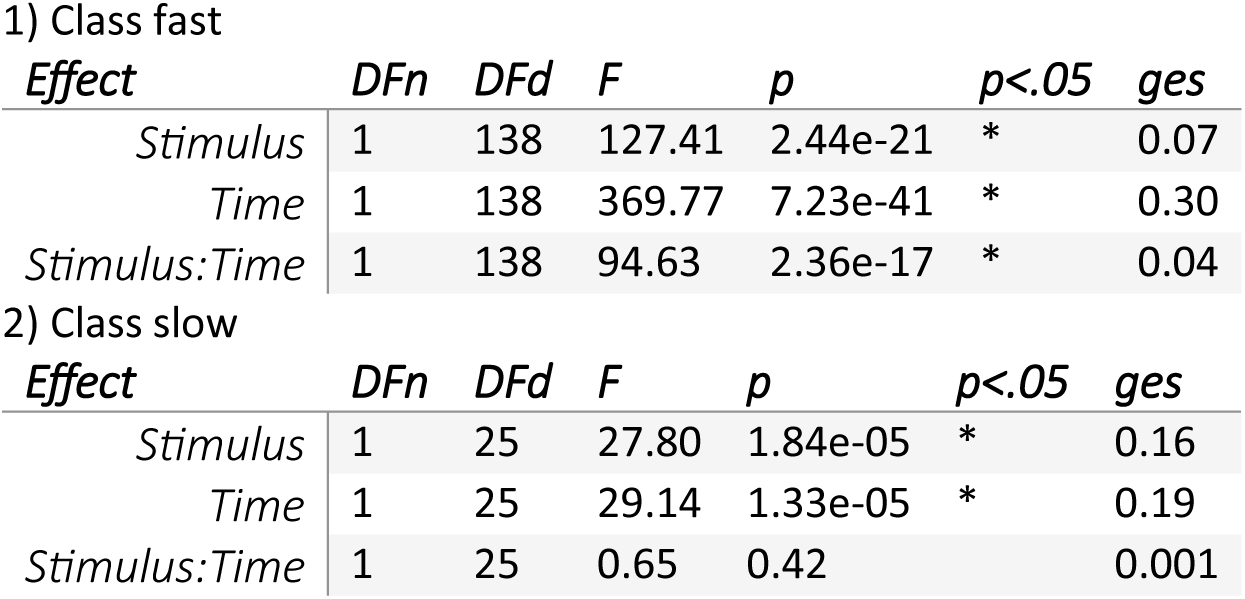
Day 2 Within-Class ANOVA.

1) Class fast
| *Effect* | *DFn* | *DFd* | *F* | *p* | *p<.05* | *ges* |
| --- | --- | --- | --- | --- | --- | --- |
| *Stimulus* | 1 | 138 | 127.41 | 2.44e-21 | * | 0.07 |
| *Time* | 1 | 138 | 369.77 | 7.23e-41 | * | 0.30 |
| *Stimulus:Time* | 1 | 138 | 94.63 | 2.36e-17 | * | 0.04 |

Supplementary Table 12: Day 2 Within-Class ANOVA
| *Effect* | *DFn* | *DFd* | *F* | *p* | *p<.05* | *ges* |
| --- | --- | --- | --- | --- | --- | --- |
| *Stimulus* | 1 | 25 | 27.80 | 1.84e-05 | * | 0.16 |
| *Time* | 1 | 25 | 29.14 | 1.33e-05 | * | 0.19 |
| *Stimulus:Time* | 1 | 25 | 0.65 | 0.42 |  | 0.001 |

**Supplementary Table 13:**
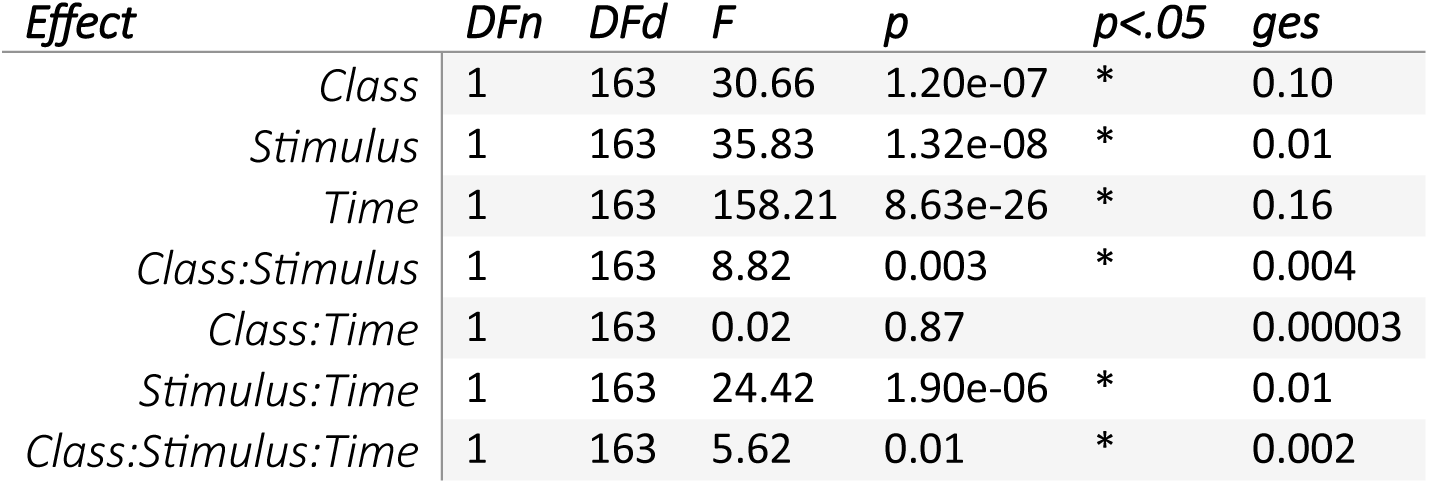
Day 3 ANOVA.

**Supplementary Table 14:** Day 3 Within-Class ANOVA.

| *Effect* | *DFn* | *DFd* | *F* | *p* | *p<.05* | *ges* |
| --- | --- | --- | --- | --- | --- | --- |
| *Stimulus* | 1 | 138 | 15.51 | 1.29e-04 | * | 0.008 |
| *Time* | 1 | 138 | 288.72 | 1.21e-35 | * | 0.26 |
| *Stimulus:Time* | 1 | 138 | 10.79 | 0.001 | * | 0.005 |

**Supplementary Table 14: Day 3 Within-Class ANOVA**
| *Effect* | *DFn* | *DFd* | *F* | *p* | *p<.05* | *ges* |
| --- | --- | --- | --- | --- | --- | --- |
| *Stimulus* | 1 | 25 | 17.16 | 0.0003 | * | 0.07 |
| *Time* | 1 | 25 | 26.15 | 2.77e-05 | * | 0.28 |
| *Stimulus:Time* | 1 | 25 | 13.68 | 0.001 | * | 0.04 |

**Supplementary Table 15:** Logistic regression results with class membership as the dependent variable and sex as predictor.

| *Predictor* | *OR* | *2.5%* | *97.5%* |
| --- | --- | --- | --- |
| *Intercept* | 0.171 | 0.103 | 0.268 |
| *Male* | 1.595 | 0.535 | 4.245 |

| *Predictor* | *Estimate* | *Std. Error* | *z value* | *p* |
| --- | --- | --- | --- | --- |
| *Intercept* | -1.766 | 0.242 | -7.300 | < .001 |
| *Male* | 0.467 | 0.520 | 0.898 | 0.369 |

### Lonsdorf et al. dataset

**Supplementary Table 16:** Day 1 ANOVA.

| *Effect* | *DFn* | *DFd* | *F* | *p* | *p<.05* | *ges* |
| --- | --- | --- | --- | --- | --- | --- |
| *Class* | 1 | 108 | 51.06 | 1.10e-10 | * | 0.19 |
| *Stimulus* | 1 | 108 | 47.68 | 3.59e-10 | * | 0.06 |
| *Time* | 1 | 108 | 28.004 | 6.40e-07 | * | 0.04 |
| *Class:Stimulus* | 1 | 108 | 0.38 | 0.53 |  | 0.0005 |
| *Class:Time* | 1 | 108 | 6.40 | 0.01 | * | 0.01 |
| *Stimulus:Time* | 1 | 108 | 8.21 | 0.005 | * | 0.008 |
| *Class:Stimulus:Time* | 1 | 108 | 6.88 | 0.009 | * | 0.007 |

**Supplementary Table 17:** Day 1 Within-Class ANOVA.

| *Effect* | *DFn* | *DFd* | *F* | *p* | *p<.05* | *ges* |
| --- | --- | --- | --- | --- | --- | --- |
| *Stimulus* | 1 | 86 | 57.23 | 3.96e-11 | * | 0.08 |
| *Time* | 1 | 86 | 75.41 | 2.16e-13 | * | 0.15 |
| *Stimulus:Time* | 1 | 86 | 0.09 | 0.75 |  | 0.0001 |

**Supplementary Table 17: Day 1 Within-Class ANOVA**
| *Effect* | *DFn* | *DFd* | *F* | *p* | *p<.05* | *ges* |
| --- | --- | --- | --- | --- | --- | --- |
| *Stimulus* | 1 | 22 | 10.64 | 0.003 | * | 0.09 |
| *Time* | 1 | 22 | 2.15 | 0.15 |  | 0.01 |
| *Stimulus:Time* | 1 | 22 | 4.45 | 0.046 | * | 0.04 |

**Supplementary Table 18:** Day 2 ANOVA.

| *Effect* | *DFn* | *DFd* | *F* | *p* | *p<.05* | *ges* |
| --- | --- | --- | --- | --- | --- | --- |
| *Class* | 1 | 108 | 25.54 | 1.77e-06 | * | 0.09 |
| *Stimulus* | 1 | 108 | 8.63 | 0.004 | * | 0.009 |
| *Time* | 1 | 108 | 95.69 | 1.45e-16 | * | 0.22 |
| *Class:Stimulus* | 1 | 108 | 0.46 | 0.49 |  | 0.0005 |
| *Class:Time* | 1 | 108 | 2.39 | 0.12 |  | 0.007 |
| *Stimulus:Time* | 1 | 108 | 0.01 | 0.90 |  | 0.00001 |
| *Class:Stimulus:Time* | 1 | 108 | 3.17 | 0.07 |  | 0.003 |

**Supplementary Table 19:** Day 2 Within-Class ANOVA.

| *Effect* | *DFn* | *DFd* | *F* | *p* | *p<.05* | *ges* |
| --- | --- | --- | --- | --- | --- | --- |
| *Stimulus* | 1 | 86 | 7.10 | 9.16e-03 | * | 0.008 |
| *Time* | 1 | 86 | 149.89 | 1.53e-20 | * | 0.37 |
| *Stimulus:Time* | 1 | 86 | 4.81 | 0.03 | * | 0.006 |

**Supplementary Table 19: Day 2 Within-Class ANOVA**
| *Effect* | *DFn* | *DFd* | *F* | *p* | *p<.05* | *ges* |
| --- | --- | --- | --- | --- | --- | --- |
| *Stimulus* | 1 | 22 | 2.64 | 0.11 |  | 0.01 |
| *Time* | 1 | 22 | 23.65 | 7.33e-05 | * | 0.20 |
| *Stimulus:Time* | 1 | 22 | 0.61 | 0.44 |  | 0.004 |

**Supplementary Table 20:** Day 3 ANOVA.

| *Effect* | *DFn* | *DFd* | *F* | *p* | *p<.05* | *ges* |
| --- | --- | --- | --- | --- | --- | --- |
| *Class* | 1 | 108 | 17.76 | 5.19e-05 | * | 0.06 |
| *Stimulus* | 1 | 108 | 3.67 | 5.78e-02 |  | 0.005 |
| *Time* | 1 | 108 | 59.59 | 6.26e-12 | * | 0.15 |
| *Class:Stimulus* | 1 | 108 | 0.30 | 0.58 |  | 0.0004 |
| *Class:Time* | 1 | 108 | 1.25 | 0.26 |  | 0.003 |
| *Stimulus:Time* | 1 | 108 | 1.85 | 0.17 |  | 0.001 |
| *Class:Stimulus:Time* | 1 | 108 | 0.05 | 0.81 |  | 0.00005 |

**Supplementary Table 21:**
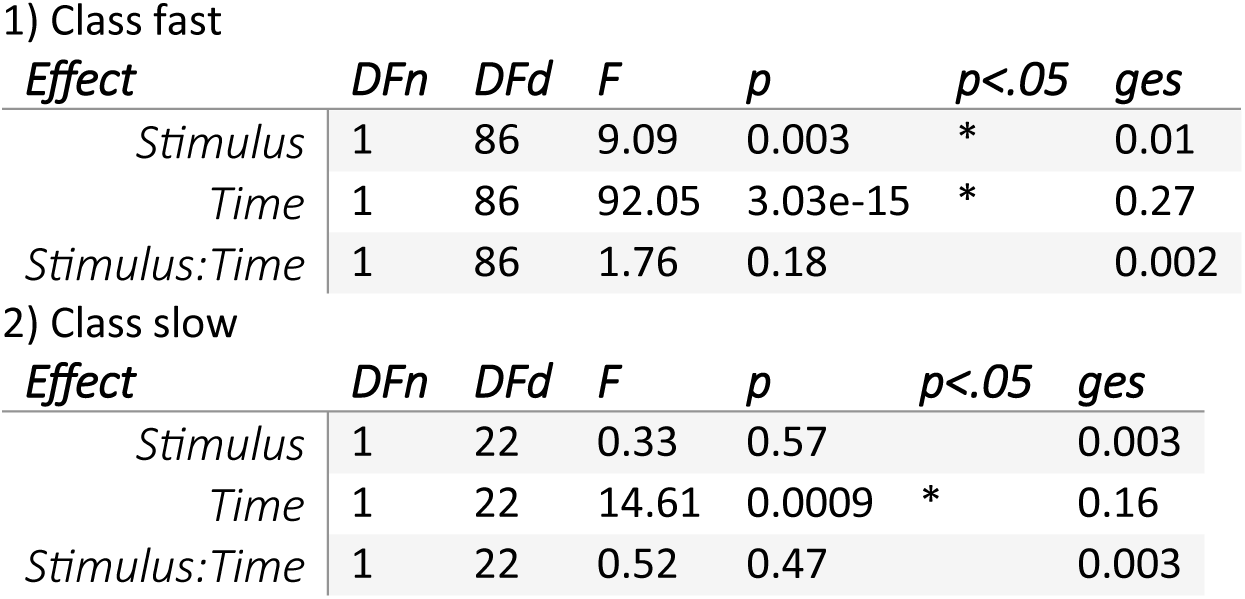
Day 3 Within-Class ANOVA.

**Supplementary Table 22:** Logistic regression results with class membership as the dependent variable and sex as predictor.

| *Predictor* | *OR* | *2.5%* | *97.5%* |
| --- | --- | --- | --- |
| *Intercept* | 0.220 | 0.116 | 0.388 |
| *Male* | 1.621 | 0.623 | 4.144 |

| *Predictor* | *Estimate* | *Std. Error* | *z value* | *p* |
| --- | --- | --- | --- | --- |
| *Intercept* | -1.513 | 0.306 | -4.937 | < .001 |
| *Male* | 0.483 | 0.479 | 1.008 | 0.313 |

